# NamZ1 and NamZ2 from the oral pathogen *Tannerella forsythia* are peptidoglycan processing exo-β-*N*-acetylmuramidases with distinct substrate specificity

**DOI:** 10.1101/2021.12.02.470905

**Authors:** Marina Borisova, Katja Balbuchta, Andrew Lovering, Alexander Titz, Christoph Mayer

## Abstract

The Gram-negative periodontal pathogen *Tannerella forsythia* is inherently auxotrophic for *N*-acetylmuramic acid (MurNAc), which is an essential carbohydrate constituent of the peptidoglycan (PGN) of the bacterial cell wall. Thus, to build up its cell wall, *T. forsythia* strictly depends on the salvage of exogenous MurNAc or sources of MurNAc, such as polymeric or fragmentary PGN, derived from cohabiting bacteria within the oral microbiome. In our effort to elucidate how *T. forsythia* satisfies its demand for MurNAc, we recognized that the organism possesses three putative orthologs of the exo-β-*N*-acetylmuramidase BsNamZ from *Bacillus subtilis*, which cleaves non-reducing end, terminal MurNAc entities from the artificial substrate pNP-MurNAc and the naturally-occurring disaccharide substrate MurNAc-β-1,4-*N*-acetylglucosamine (GlcNAc). TfNamZ1 and TfNamZ2 were successfully purified as soluble, pure recombinant His_6_-fusions and characterized as exo-lytic β-*N*-acetylmuramidases with distinct substrate specificities. The activity of TfNamZ1 was considerably lower compared to TfNamZ2 and BsNamZ, in the cleavage of pNP-MurNAc and MurNAc-GlcNAc. When peptide-free PGN glycans were used as substrates, we revealed striking differences in the specificity and mode of action of these enzymes, as analyzed by mass spectrometry. TfNamZ1, but not TfNamZ2 or BsNamZ, released GlcNAc-MurNAc disaccharides from these glycans. In addition, glucosamine (GlcN)-MurNAc disaccharides were generated when partially N-deacetylated PGN glycans from *B. subtilis 168* were applied. This characterizes TfNamZ1 as a unique disaccharide-forming exo-lytic β-*N*-acetylmuramidase (exo-disaccharidase), and, TfNamZ2 and BsNamZ as sole MurNAc monosaccharide-lytic exo-β-*N*-acetylmuramidases.

**IMPORTANCE:** Two exo-β-*N*-acetylmuramidases from *T. forsythia* belonging to glycosidase family GH171 (www.cazy.org) were shown to differ in their activities, thus revealing a functional diversity within this family: NamZ1 releases disaccharides (GlcNAc-MurNAc/GlcN-MurNAc) from the non-reducing ends of PGN glycans, whereas NamZ2 releases terminal MurNAc monosaccharides. This work provides a better understanding of how *T. forsythia* may acquire the essential growth factor MurNAc by the salvage of PGN from cohabiting bacteria in the oral microbiome, which may pave avenues for the development of anti-periodontal drugs. On a broad scale, our study indicates that the utilization of PGN as a nutrient source, involving exo-lytic *N*-acetylmuramidases with different modes of action, appears to be a general feature of bacteria, particularly among the phylum Bacteroidetes.

## INTRODUCTION

The net-shaped heteropolymer peptidoglycan (PGN) is the major structural component of the cell wall of bacteria, conferring physical strength to the cells. It is composed of β-1,4-linked glycan chains of alternating *N*-acetylmuramic acid (MurNAc) and *N*-acetylglucosamine (GlcNAc) monosaccharides, which are interconnected via oligopeptides attached to MurNAc [Walter, 2019 #19888]. The PGN cell wall is continuously degraded and remodelled during bacterial growth and differentiation (i.e., PGN turnover), a process that involves the combined activity of potentially autolytic enzymes (autolysins), including endo-cleaving *N*-acetylmuramidases, *N*-acetylglucosaminidases, and lytic transglycosidases [Oshida, 1995 #15478;Mayer, 2012 #13163;Dik, 2017 #16124]. Since the synthesis and integrity of the PGN is essential for bacteria to cope with environmental stresses, antibacterial-acting endo-lytic PGN glycosidases that ultimately cause bacterial lysis, such as endo-*N*-acetylmuramidases, commonly known as lysozymes, represent basal factors of innate immunity against bacterial infections [Ragland, 2017 #19745].

The process of PGN turnover and utilization as a nutrient and energy source (i.e., PGN recycling or salvage), also involves the activity of exo-lytic hydrolases, which processively degrade PGN from the ends, thereby averting uncontrolled autolysis [Litzinger, 2010 #12434]. We previously showed that *Bacillus subtilis* is able to digest intact PGN by sequential hydrolysis from the non-reducing ends via the joined activity of the exo-β-*N*-acetylglucosamidase BsNagZ, the exo-*N*-acetylmuramoyl-L-alanine amidase AmiE, and the exo-lytic *N*-acetylmuramidase BsNamZ, which sequentially release GlcNAc, peptides, and MurNAc, respectively [Litzinger, 2010 #12434; Müller, 2021 #18933]. Presumably, these hydrolases are involved in the degradation of PGN during turnover of the cell wall or the decay of lysed cells in a starving population of *B. subtilis* [Borisova, 2016 #14239].

The exo-lytic β-*N*-acetylmuramidase BsNamZ, was recently characterized and shown to constitute a novel family of glycosidases (CAZy GH171; www.cazy.org/GH171.html) [Müller, 2021 #18933]. BsNamZ uniquely cleaves terminal *N*-acetylmuramic acid (MurNAc) moieties from the non-reducing ends (exo-lytic cleavage) of peptide-free, PGN glycans and also hydrolyzes the artificial substrate 4-nitrophenyl β-MurNAc (pNP-MurNAc). Thus, the activity of BsNamZ differs from lysozyme-like endo-*N*-acetylmuramidases (CAZy GH25), which are unable to cleave terminal MurNAc residues but hydrolyze internal glycosidic bonds within the PGN network (endo-lytic cleavage) and generally require peptide-substituted MurNAc residues [Ragland, 2017 #19745].

We previously recognized that putative NamZ orthologs are narrowly distributed within bacteria and are frequently found within members of the Bacteroidetes phylum [Müller, 2021 #18933]. The oral pathogen *Tannerella forsythia*, which is strongly associated with disease, such as periodontitis and rheumatoid arthritis, belongs to the Bacteroidetes [Loyola-Rodriguez, 2010 #19862;Martinez-Rivera, 2017 #19873;Bourgeois, 2019 #19864]. Intriguingly, this organism is a natural auxotroph for MurNAc [Wyss, 1989 #9982]. Since *T. forsythia* constitutes a classical Gram-negative PGN cell wall [Mayer, 2019 #16115], it strictly depends on a supply of MurNAc, for growth in axenic culture, or sources of MurNAc, which may be provided by other bacteria, to proliferate in the oral habitat [Wyss, 1989 #9982;Hottmann, 2018 #14525;Hottmann, 2021 #19871]. Salvage of MurNAc from the medium, requires the MurNAc-specific inner membrane transporter TfMurT of *T. forsythia* [Ruscitto, 2016 #14240]. In addition, *T. forsythia* imports PGN-derived disaccharides, particularly GlcNAc-MurNAc and GlcNAc-1,6-anhydroMurNAc, and possibly also muropeptides, via the recently identified transporter TfAmpG [Ruscitto, 2018 #16099;Mayer, 2020 #18932]. The latter transporter was recently shown to be is required for the utilization of polymeric PGN as a nutrient source by *T. forsythia* [Mayer, 2020 #18932]. However, so far, the *T. forsythia* enzymes responsible for the release of MurNAc and MurNAc-containing disaccharides from PGN have remained unknown.

*T. forsythia* is able to utilize polymeric or fragmentary PGN to satisfy its need for MurNAc [Ruscitto, 2017 #16112;Ruscitto, 2018 #16099;Mayer, 2020 #18932;Hottmann, 2021 #19871]. Intriguingly, we recognized that *T. forsythia* harbours three putative *namZ* orthologs on its chromosome, which we named TfNamZ1-3 [Hottmann, 2021 #19871]. It is reasonable to assume that these NamZ enzymes may be critical for the organism to liberate the essential growth factor MurNAc, however, likely they do not all serve the same purpose. Thus we aimed to elucidate the activities and substrate specificities of the NamZ orthologs from *T. forsythia*. Here we present the characterization of two of them and show that, despite of their high amino acid sequence identity, TfNamZ1 and TfNamZ2 differ in their substrate specificity and product formation. TfNamZ1 is an exo-β-*N*-acetylmuramidase that releases primarily GlcNAc-MurNAc from the non-reducing end of peptide-free PGN glycan strands, whereas TfNamZ2 releases MurNAc from the non-reducing ends of the glycans and thus, possesses classical exo-β-*N*-acetylmuramidase activity, similar to that previously described for BsNamZ [Müller, 2021 #18933]. Interestingly, TfNamZ1 is encoded within a putative operon in the *T. forsythia* genome that also contains the genes encoding TfMurT and TfAmpG, as well as a number of other enzymes related to PGN salvage (Fig. 1). Thus, TfNamZ1 and TfNamZ2, yield the substrates of the transporters TfAmpG and TfMurT, respectively, thereby satisfying *T. forsythia*’s need for the crucial growth factor MurNAc.

**FIG. 1.**
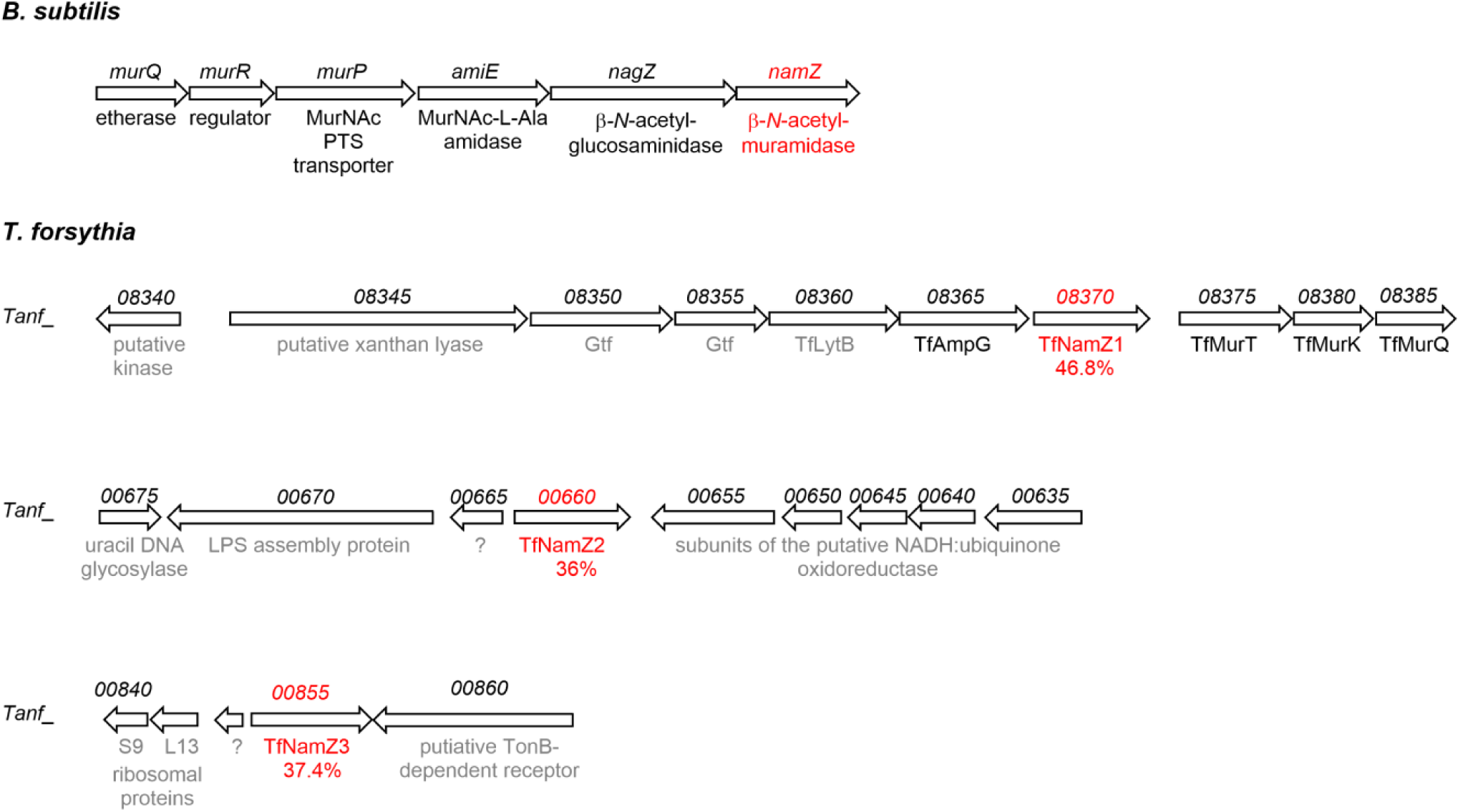
Genomic organization of the exo-*N*-acetylmuramidase *namZ* of *B. subtilis* and the three putative *namZ*-like genes of *T. forsythia*. The *namZ* operon of *B. subtilis* strain 168 (NCBI Reference Sequence accession no. NC_000964.3) includes three genes (*amiE*, *nagZ* and *namZ*) involved in PGN turnover/recycling and three genes (*murQRP*) required for the uptake of MurNAc and the transcriptional regulation of the operon: *amiE* encodes the exo-MurNAc-L-Ala amidase BsAmiE [Litzinger, 2010 #12434], *nagZ* encodes the exo-β-*N*-acetylglucosaminidase BsNagZ [Litzinger, 2010 #12680] and *namZ* encodes the recently identified and characterized exo-β-*N*-acetylglucosaminidase BsNamZ (in red) [Müller, 2021 #18933]; furthermore, *murP* encodes the MurNAc-specific phosphotransferase system (PTS) transporter BsMurP, *murQ* encodes the MurNAc 6-phosphate etherase MurQ and *murR* encodes the putative transcriptional regulator of the operon [Borisova, 2016 #14239]. Within the genome of *T. forsythia* strain ATCC 43037 (NCBI Reference Sequence: NZ_JUET00000000.1), three orthologous *namZ*-like family GH171 glycosidases (CAZy GH171; www.cazy.org) were identified, *TfnamZ1 (Tanf_08370)*, *TfnamZ2 (Tanf_00660)* and *TfnamZ3* (*Tanf_00855*). According to the Kegg Database [Kanehisa, 2000 #20023], the *TfnamZ1* cluster of *T. forsythia* includes 9 genes (*Tanf_08345* to *Tanf_08385*), with *Tanf*_*08375, Tanf_08380 and Tanf_08385,* encoding the MurNAc transporter TfMurT [Ruscitto, 2016 #14240], the MurNAc kinase TfMurK [Hottmann, 2018 #14525], and the MurNAc 6-phosphate etherase TfMurQ [Ruscitto, 2016 #14240], respectively, which are involved in MurNAc transport and catabolism. The genes upstream of *TfnamZ1 (Tanf_08370)* encode the inner membrane permease TfAmpG *(Tanf_08365)*, which was recently shown to transport the disaccharides GlcNAc-MurNAc and GlcNAc-anhMurNAc [Mayer, 2020 #18932] and a putative lytic transglycosylase, which we named TfLytB (*Tanf_08360)*. The functions of the further upstream genes are unknown: *Tanf_08350* and *Tanf_08355* encode two putative glycosyltransferases (Gtf), and *Tanf_08345* a putative xanthan lyase. The gene *TfnamZ2 (Tanf_00660)* is organized in a gene cluster with three other genes, *Tanf_00675*, *Tanf_00670* and *Tanf_00665*, which encode an uracil DNA glycosylase, LPS assembly protein and a hypothetical protein, respectively. The third *namZ* ortholog of *T. forsythia*, *TfnamZ3* (*Tanf_00855*) is a monocistronic gene. Genes that code for proteins of a yet uncharacterized function, indicated by a question mark, are presented in gray, characterized proteins are shown in black, and the *namZ* genes are shown in red.

## RESULTS

### Identification of three putative NamZ orthologs from *T. forsythia*

A BLAST search, using the amino acid sequence of the exo-β-*N*-acetylmuramidase BsNamZ [Müller, 2021 #18933], identified three putative orthologs (DUF1343 family proteins; http://pfam.xfam.org/) in the *T. forsythia* ATCC 43037 proteome, which we named TfNamZ1 (Tanf_08370), TfNamZ2 (Tanf_00660), and TfNamZ3 (Tanf_00855). They display overall amino acid sequence identities of 46.8%, 36.0% and 37.4%, respectively, to BsNamZ [Hottmann, 2021 #19871]. In addition, the BLAST search revealed two NamZ-like DUF1343 family proteins from the phylogenetically related *Bacteroides fragilis* strain NCTC 9343 (BF9343_0362 and BF9343_0369), for which high resolution crystal structures have been determined (www.rcsb.org; PDB-IDs: 4K05 and 4JJA). These proteins, named BfNamZ1 and BfNamZ2, share 43.0% and 41.2% overall amino acid identity with BsNamZ, respectively. A multiple amino acid sequence alignment and a phylogenetic tree of the different putative NamZ orthologs from *T. forsythia, B. fragilis* and *B. subtilis* is depicted in Figure S1 (see Supplemental Material). The phylogenetic tree revealed that TfNamZ1, and BfNamZ1 form a branch and a second branch includes BfNamZ2 and the *T. forsythia* orthologs TfNamZ2 and TfNamZ3. This indicated BsNamZ-related functions of TfNamZ1 and BfNamZ2, and potentially somewhat distinct functions of TfNamZ2, TfNamZ3 and BfNamZ2, although prediction of variations in function is difficult solely on the basis of primary amino acid sequence.

To obtain some hints regarding their function, we analyzed the genomic organization of the three NamZ-encoding genes of *T. forsythia* (Fig. 1). *TfnamZ1* is located within a large putative operon attributed to the catabolism/recycling of MurNAc and PGN-derived disaccharides, between the coding gene of the recently characterized GlcNAc-MurNAc/GlcNAc-anhMurNAc transporter TfAmpG [Ruscitto, 2017 #16112; Mayer, 2020 #18932] and the three genes encoding the MurNAc-transporter TfMurT [Ruscitto, 2016 #14240], the MurNAc kinase TfMurK [Hottmann, 2018 #14525], and the MurNAc 6-phosphate lactyl ether hydrolase TfMurQ [Ruscitto, 2016 #14240]. Thus, the genomic organization of TfNamZ1 indicated a function related to PGN catabolism and suggested that TfNamZ1 may play a role in the formation of substrates of the TfMurT and TfAmpG transporters [Hottmann, 2021 #19871]. In contrast, the genomic context of *TfnamZ2* and *TfnamZ3* did not allow any conclusions on their cellular function. Interestingly, the coding genes of the two NamZs from *B. fragilis* are both located within a gene cluster that resembles the *TfnamZ1* operon (cf. Fig. 1) and, thus, likely play a role in PGN recycling/catabolism as well.

### TfNamZ1 and TfNamZ2 are exo-β-*N*-acetylmuramidases that hydrolyse pNP-MurNAc and MurNAc-GlcNAc

To compare their activities, substrate specificities and biochemical functions, we cloned and heterologously overexpressed the three TfNamZ enzymes recombinantly. A first attempt to purify the recombinant proteins failed, due to the insolubility of the full length forms (data not shown). However, with the aid of the multiple amino acid sequence alignment (Fig. S1) and signal peptide predictions (Signal P 5.0), we realized that all three TfNamZs proteins contain putative signal sequences for protein secretion. We thus constructed recombinant proteins that lack the putative signal sequences (see Table S1 for a full list of the used primers) and this led to successful expression of the recombinant C-terminal His_6_ fusion proteins (Fig. S2). Subsequently, TfNamZ1 and TfNamZ2 were purified to apparent homogeneity using Ni-affinity chromatography and gel filtration, as demonstrated by SDS-PAGE (Fig. S2). The molecular weights of these proteins were in agreement with the expected size of 45.8 kDa and 43.3 kDa, respectively. The protein yields from three independent purifications were in the range of 20 to 23 mg/L for TfNamZ1 and TfNamZ2. For so far unknown reasons, we failed to isolate TfNamZ3, either with or without putative signal sequence. Thus, we restricted our studies to the functional characterization of TfNamZ1 and TfNamZ2, but excluded TfNamZ3 from further biochemical investigation.

The exo-β-*N*-acetylmuramidase BsNamZ has previously been shown to hydrolyse solely the chemically synthesized, chromogenic substrate pNP-MurNAc, whereas the exo-β-N-acetylglucosaminidase BsNagZ hydrolyses exclusively pNP-GlcNAc [Litzinger, 2010 #12434; Litzinger, 2010 #12680;Müller, 2021 #18933]. We first analyzed whether the recombinant TfNamZ1 and TfNamZ2 proteins also specifically cleave pNP-MurNAc. This was indeed the case, as visualized by the development of the yellow colored reaction product, i.e., the release of para-nitrophenol from pNP-MurNAc (Fig. 2). Both *T. forsythia* enzymes did not cleave pNP-GlcNAc, which indicates that they are specific exo-β-*N*-acetylmuramidases, such as BsNamZ (Fig. 2). Notably, the reaction of TfNamZ1 with pNP-MurNAc was considerably slower as compared to TfNamZ2 and BsNamZ when identical enzyme concentrations were applied, which indicates that TfNamZ1 is a less efficient exo-β-*N*-acetylmuramidase for the cleavage of the chromogenic pNP-MurNAc substrate.

**FIG. 2.**
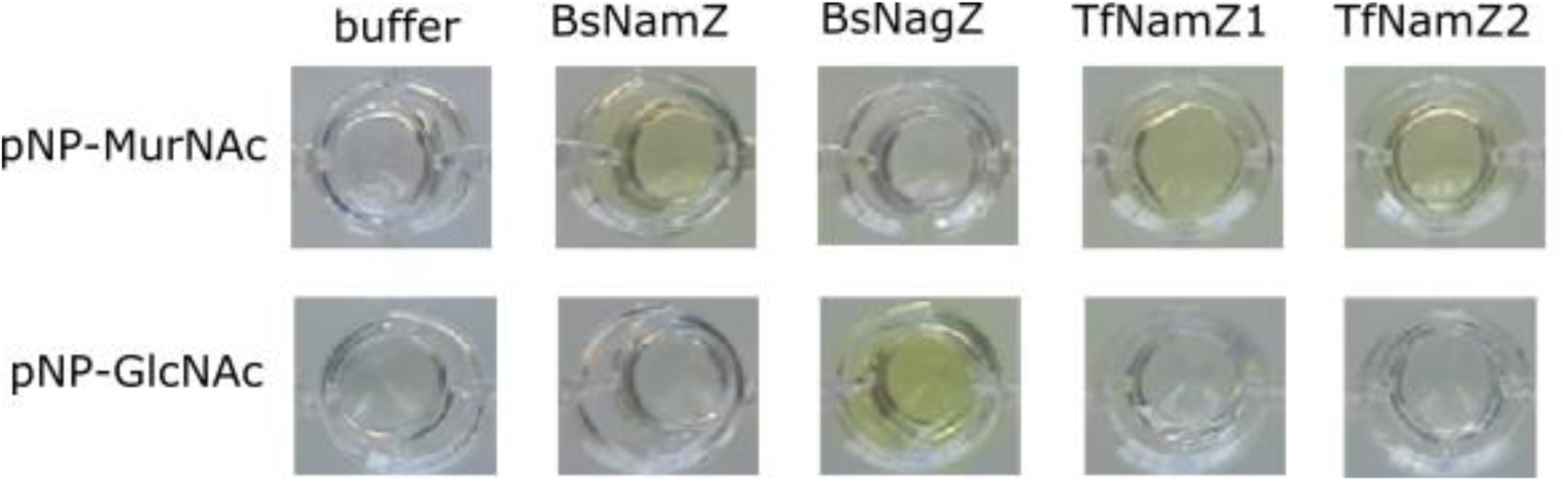
Specificity of NamZ proteins for the chromogenic substrates pNP-GlcNAc and pNP-MurNAc. The recombinant enzymes TfNamZ1 and TfNamZ2 were assayed, together with two recombinant enzymes from *B. subtilis* 168 of known activity. The β-*N*-acetylmuramidase BsNamZ cleaves pNP-MurNAc and the β-*N*-acetylglucosaminidase BsNagZ cleaves pNP-GlcNAc, as evidenced by the development of a yellow colour upon the release of para-nitrophenol. TfNamZ1 and TfNamZ2, both showed exo-β-*N*-acetylmuramidase activity, i.e., they specifically cleaved pNP-MurNAc, and showed no β-*N*-acetylglucosaminidase activity, i.e. no cleavage of pNP-GlcNAc.

We next analyzed whether TfNamZ1 and TfNamZ2 can also cleave the PGN-derived disaccharide MurNAc-GlcNAc, which constitutes the minimal natural substrate of the exo-β-*N*-acetylmuramidase BsNamZ [Müller, 2021 #18933]. MurNAc-GlcNAc was obtained by enzymatic digestion of PGN from *S. aureus* with recombinant Atl *N*-acetylmuramyl-L-Ala amidase SaAtl^AM^ and recombinant Atl *N*-acetylglucosaminidase SaAtl^Glc^ [Oshida, 1995 #15478;Nega, 2020 #19874]. LC-MS analyses revealed that Atl releases not only the disaccharide MurNAc-GlcNAc, which is the main carbohydrate product, as recently described [Nega, 2020 #19874], but also the trisaccharide GlcNAc-MurNAc-GlcNAc in about 15 times lower abundance (data not shown). To increase the amount of MurNAc-GlcNAc in the preparation, we further digested the trisaccharide containing Atl-hydrolysate with the exo-β-*N*-acetylglucosaminidase BsNagZ, thereby generating MurNAc-GlcNAc and GlcNAc. This MurNAc-GlcNAc preparation was then used as a substrate to test the activity of TfNamZ1 and TfNamZ2 by LC-MS analysis. The exact monoisotopic masses of the proton adducts ([M+H]^+^) of the investigated PGN metabolites MurNAc-GlcNAc (substrate), MurNAc and GlcNAc (products) are listed in Table S3. The MurNAc-GlcNAc preparation contained only minor amounts of MurNAc and GlcNAc prior to enzymatic incubation (Fig. 3). The incubation with TfNamZ2 and BsNamZ overnight at 37 °C led to a complete degradation of the disaccharide and the respective formation of the monosaccharides MurNAc and GlcNAc (Fig. 3). Surprisingly, however, when the same amount of TfNamZ1 was added, only about half of the MurNAc-GlcNAc substrate was digested and consequently also only about half of the amounts of the products were generated (Fig. 3). Thus, TfNamZ1 has considerably lower activity with both substrates, MurNAc-GlcNAc and pNP-MurNAc. Neither TfNamZ1 nor TfNamZ2 were able to hydrolyze GlcNAc-MurNAc, as shown by LC-MS analysis (Fig. S3). In contrast, the disaccharide GlcNAc-MurNAc was completely degraded by the exo-*N*-acetylglucosaminidase BsNagZ, yielding GlcNAc and MurNAc (Fig. S3).

**FIG. 3.**
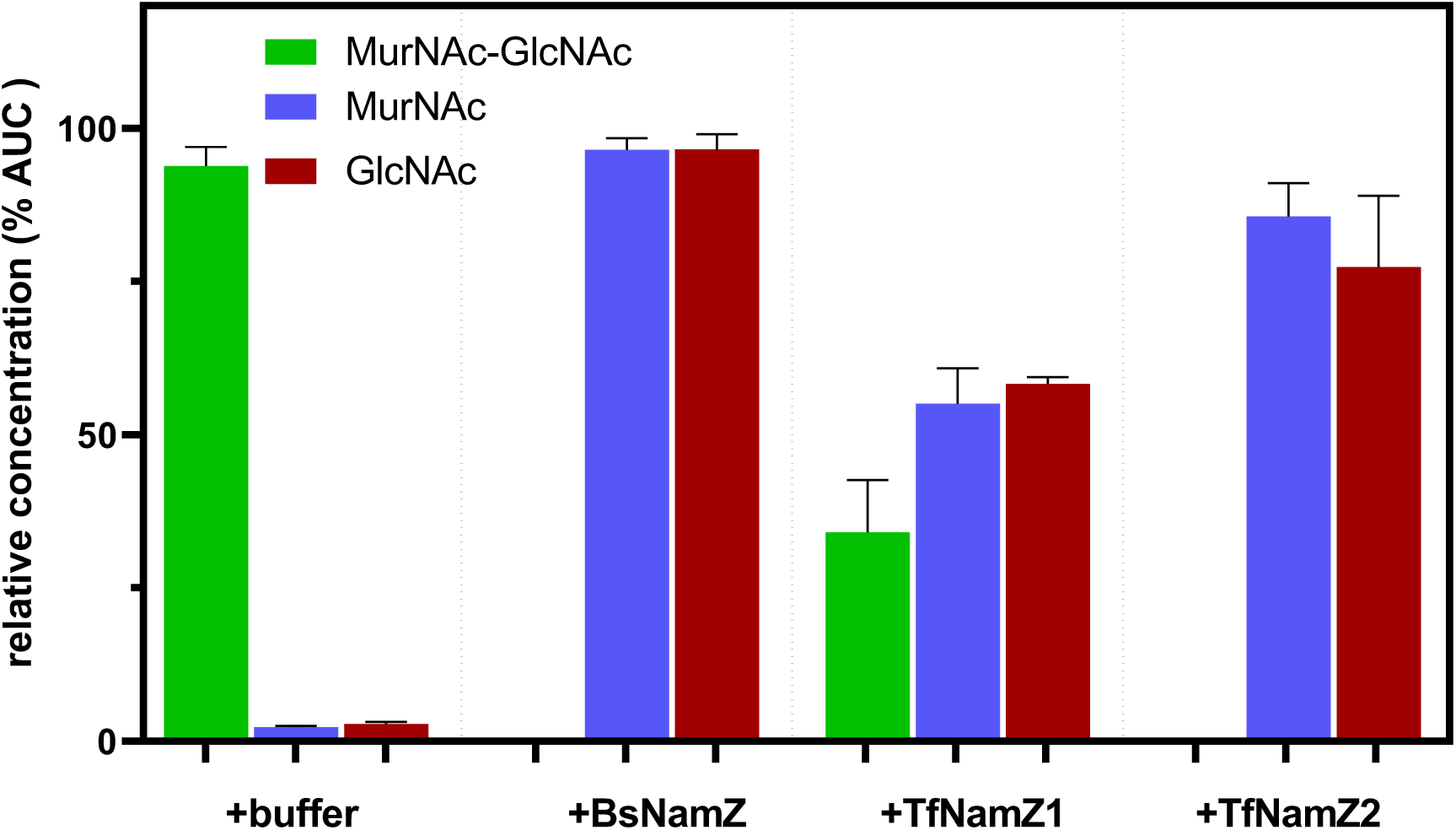
Cleavage of MurNAc-GlcNAc by BsNamZ, TfNamZ1 and TfNamZ2. MurNAc-GlcNAc, generated by digestion of *S. aureus* peptidoglycan (PGN) with SaAtl^AM^, *Sa*Atl^Glc^ and BsNagZ was incubated for 20 h at 37 °C with the BsNamZ, TfNamZ1, TfNamZ2 or buffer (as control). The remaining substrate (MurNAc-GlcNAc, green) in the reaction mixture and the reaction products (MurNAc, blue; GlcNAc, red) were quantified using LC-MS. The relative metabolite concentrations (% area under the curve, AUC, of the respective mass peak) represent the mean from three biological replicates.

### TfNamZ1 releases GlcNAc-MurNAc disaccharides from peptide-free PGN glycans

Differences in the activities of the two TfNamZs with the pNP-MurNAc and MurNAc-GlcNAc indicated that the proteins may have distinct substrate preferences. Therefore, we compared the activities of TfNamZ1 and TfNamZ2 using peptide-free PGN glycans. For this purpose, we generated short strands of peptide-free PGN glycans with varying length by digestion of *S. aureus* PGN with the Atl^AM^ amidase, as described [Oshida, 1995 #15478; Nega, 2020 #19874]. When TfNamZ1 was added to the *S. aureus*- derived glycan strands, a main product with [M+H]^+^ = 497.198 m/z, corresponding to a disaccharide containing the sugars GlcNAc and MurNAc, was formed (Fig. 4A). We identified the disaccharide as GlcNAc-MurNAc, through the degradation with the exo-β-*N*-acetylglucosaminidase BsNagZ, yielding GlcNAc and MurNAc (Fig. 4A and 4B). In contrast, when TfNamZ2 was incubated with PGN glycans, we observed no release of disaccharides. Only small amounts of MurNAc were released, presumably due to the cleavage of terminal MurNAc residues from the non-reducing ends occasionally occurring in the PGN glycan strands (Figure 4B). Also upon incubation with TfNamZ1 small amounts of MurNAc were detected, which is an agreement with the above function of TfNamZ1 in hydrolyzing nonreducing-end terminal MurNAc residues from pNP-MurNAc and MurNAc-GlcNAc.

**FIG. 4.**
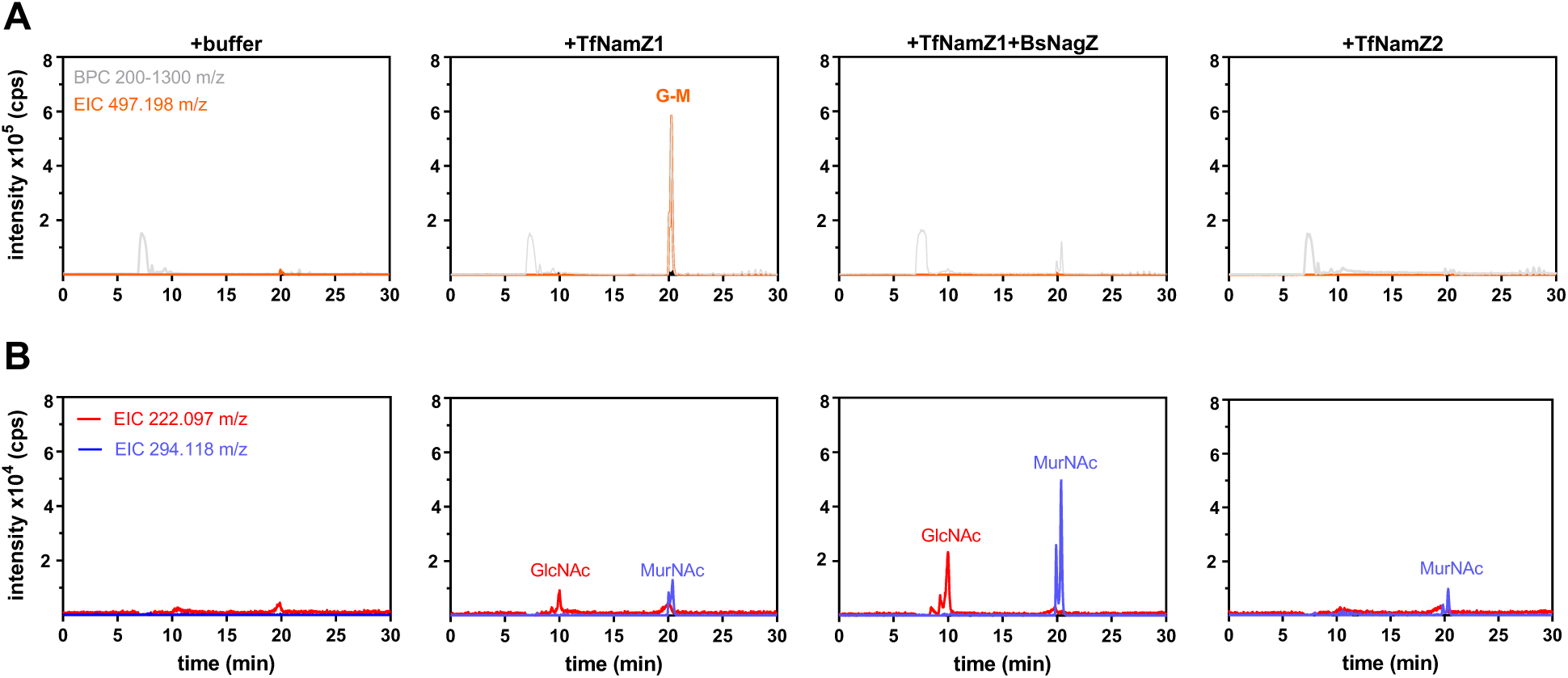
TfNamZ1 and TfNamZ2 show differences in product formation when incubated with PGN-derived glycans from *S. aureus*. Peptide-free PGN glycans were generated by digestion of PGN sacculi isolated from *S. aureus* strain USA300 with the *S. aureus* amidase SaAtl^AM^. The PGN glycan strands were incubated for 20 hrs at 37 °C with TfNamZ1 or TfNamZ2 recombinant enzymes, or buffer as a control (as indicated). Subsequently, the TfNamZ1 digested glycan strands were further digested with BsNagZ (TfNamZ1+BsNagZ). The metabolite concentrations (peak intensities; cps) in the reaction mixtures were analyzed by LC-MS. **A.** Base peak chromatograms (gray lines) and extracted ion chromatograms (EIC) corresponding to GlcNAc-MurNAc (orange), with [M+H]^+^ = 497.198 m/z are presented. **B.** EICs, corresponding to GlcNAc (red) and MurNAc (blue), with [M+H]^+^ = 222.097 m/z and [M+H]^+^ = 294.118 m/z, respectively, were plotted.

Notably, small amounts of GlcNAc were identified in the TfNamZ1-digest of PGN glycans but not with TfNamZ2 (Fig. 4B), which likely results from the release of GlcNAc residues located at the reducing end of the glycan strands by TfNamZ1 (Fig. 4B). In agreement with this assumption, TfNamZ1, but not TfNamZ2, cleaved the trisaccharide GlcNAc-MurNAc-GlcNAc ([M+H]^+^ =700.277 m/z), which is present in minor amounts in the *S. aureus* PGN glycan strands preparation (Fig. S4A). Consistently, the tetrasaccharide MurNAc-GlcNAc-MurNAc-GlcNAc, which is also present in low amounts in this preparation, was cleaved twice by TfNamZ1, releasing MurNAc, GlcNAc-MurNAc and GlcNAc, whereas this substrate was cleaved only once by TfNamZ2, yielding MurNAc and GlcNAc-MurNAc-GlcNAc residues (Fig. S4A and S4B, see also Fig. 4).

To better characterize the substrate specificity of the TfNamZ proteins, we generated peptide-free, PGN glycan strands also from *B. subtilis*, by digestion of whole PGN with the MurNAc-L-Ala amidase CwlC of *B. subtilis* as reported [Müller, 2021 #18933]. As expected, glycan strands from *B. subtilis* were not substrates for the TfNamZ2 enzyme (Fig. S5). However, with this substrate the unique disaccharide-releasing activity of the exo-β-*N*-acetylmuramidase TfNamZ1 was confirmed. The digestion of PGN glycans from *B. subtilis* by TfNamZ1 yielded a product with mass of [M+H]^+^ = 497.198 m/z, corresponding to GlcNAc-MurNAc (Fig. 5A), which in turn could be digested by BsNagZ, yielding GlcNAc and MurNAc, but was not cleaved by BsNamZ (Figs. 5A and 5B). Interestingly, we could also show that TfNamZ1 also generates a further product with the mass [M+H]^+^ = 455.188 m/z (Fig. 5A). This mass corresponds to a disaccharide containing *N*-acetylglucosamine GlcN and MurNAc, likely GlcN-MurNAc, since it was neither cleaved by BsNagZ nor by BsNamZ (Fig.5A). Apparently, TfNamZ1 displays a rather broad specificity with respect to the sugar moiety at the nonreducing end, which may be GlcNAc or GlcN.

**FIG. 5.**
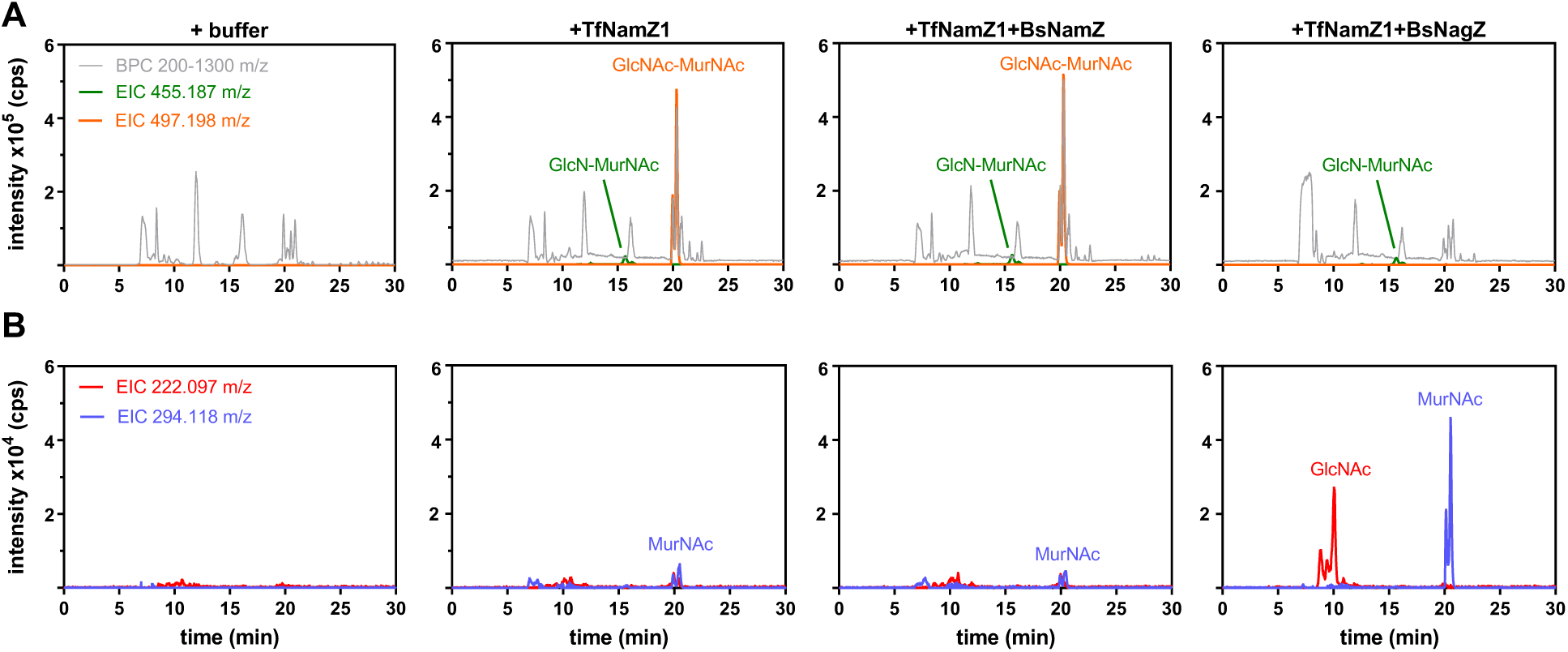
Substrate specificity of TfNamZ1 with PGN-derived glycans from *B. subtilis.* PGN glycan strands from *B. subtilis* were generated by digestion of PGN sacculi isolated from *B. subtilis* 168 with the *B. subtilis* amidase BsCwlC. The enzyme reactions were heat inactivated and the generated *B. subtilis* glycan strands were incubated with TfNamZ1 or buffer as a control. Subsequently, the TfNamZ1 digested glycan strands were further digested with BsNamZ (TfNamZ1+BsNamZ) or with BsNagZ (TfNamZ1+BsNagZ). Enzyme reactions were analyzed by LC-MS in positive ion mode. **A**. Data are presented as base peak chromatograms 200-1300 m/z (gray lines) and as extracted ion chromatogram (EIC), corresponding to GlcNAc-MurNAc (orange) and GlcN-MurNAc (green) with [M+H]^+^ = 497.198 m/z and or of [M+H]^+^ = 455.187 m/z, respectively, are shown as intensities (cps). **B**. Data are presented as EICs (intensity; cps), corresponding to GlcNAc (red) and MurNAc (blue), with [M+H]^+^ = 222.097 m/z (red) and [M+H]^+^ = 294.118 m/z (blue), respectively.

We could further show that intact *B. subtilis* PGN sacculi, which contain glycans harbouring peptide stems and crosslinks, are not substrates for TfNamZ1 and TfNamZ2 (Fig. S6). Only small amounts of GlcNAc-MurNAc disaccharides were released from *B. subtilis* PGN sacculi with TfNamZ1, which presumably derived from small parts of the PGN sacculi that lack peptide stems. No peptide-containing disaccharides or MurNAc-peptides were identified by LC-MS as potential products of TfNamZ1 or TfNamZ2. It should be noted that the amidase CwlC removes both, cross-linked and non-cross-linked peptides from the *B. subtilis* sacculi (Fig. S6). The peptide-free PGN glycan polysaccharide chains are not detectable by LC-MS due to their size, however these are readily hydrolysed by TfNamZ1 and the reaction product, which is mainly GlcNAc-MurNAc (besides GlcN-MurNAc, in the case of applying in PGN preparations from *B. subtilis*) can be detected as shown above.

## DISCUSSION

This study showed that TfNamZ1 (Tanf_08370) and TfNamZ2 (Tanf_00660) of *T. forsythia,* two members of family 171 of glycosidases (CAZy GH171; www.cazy.org), are exo-lytic β-*N*-acetylmuramidase with distinct substrate specificity and product outcome. They revealed distinct activities when peptide-free PGN glycans were used as substrates. They, hence constitute different subgroups within the family 171 of glycosidases. TfNamZ2, like BsNamZ, only cleaved terminal MurNAc entities from the non-reducing-ends of the PGN glycans, whereas TfNamZ1 primarily released GlcNAc-MurNAc and GlcN-MurNAc disaccharides from PGN glycans (Figs. 4 and 6). Although, both enzymes were able to hydrolyze pNP-MurNAc and MurNAc-GlcNAc, only TfNamZ2 showed an activity similar to BsNamZ, the previously characterized prototypic exo-β-*N*-acetylmuramidase and founding member of this glycosidase family [Müller, 2021 #18933]. TfNamZ1 had an about 50% lower activity compared to TfNamZ2 and BsNamZ, when MurNAc-GlcNAc and pNP-MurNAc were used as substrates. It may be speculated that TfNamZ1 and TfNamZ2 differ in their modes of substrate binding, in that the former allows accommodation of a non-reducing terminal disaccharide (GlcNAc-MurNAc/GlcN-MurNAc), whereas the latter enzyme only accommodates a non-reducing terminal MurNAc monosaccharide, possibly with higher affinity which could explain the higher enzymatic activity of TfNamZ2 (and BsNamZ) with MurNAc-glycoside substrates. Further structure-function studies are required to fully appreciate the differences in the substrate preferences within the different NamZ enzymes. The different mode of cleavage of TfNamZ1 was unexpected, since the protein share high overall amino acid sequence identity with the exo-β-*N*-acetylmuramidase NamZ from *B. subtilis* (46.8%), whereas the functionally analogous TfNamZ2 share only 36% identity (and TfNamZ3 37.4%). This indicates that predictions of the substrate specificities of NamZs cannot be achieved solely on the basis of the primary structure of the proteins (Fig. S1). Accordingly, BsNamZ displays 43.0% amino acid sequence identity with BfNamZ1 and 41.2% with BfNamZ2, however BfNamZ1 clusters with TfNamZ1 in a neighbour-joining tree (Fig. S1), suggesting that they are functionally related. Hence, further studies exploring the structure-function of the TfNamZ glycosidases will clarify differences in their catalytic center and structural determinants for the two modes of exo-lytic activities, as well as elucidate the mechanism of action of this family of glycosidases. Clearly, both TfNamZs require substrates, in which the MurNAc carries a free carboxylic acid at the lactyl group that is not substituted by a peptide stem. In agreement with this, we previously showed that a methyl ester derivative of pNP-MurNAc did not serve as a substrate for BsNamZ [Müller, 2021 #18933]. It is tempting to speculate that this carboxylic acid is crucial for specific binding of MurNAc and possibly directly involved in glycoside cleavage.

We recently proposed a pathway whereby *T. forsythia* salvages exogenous PGN derived from the cohabiting bacteria within the oral microbial community for the provision of the essential growth factor MurNAc [Mayer, 2020 #18932; Hottmann, 2021 #19871]. This pathway involves PGN uptake and the concomitant removal of its stem peptides by a so far unidentified MurNAc-L-Ala amidase. The removal of the stem peptides of the exogenous PGN may be used by *T. forsythia* as a mechanism to discriminate its own PGN and thus protects it from enzymatic degradation [Hottmann, 2021 #19871]. Peptide-free PGN glycan strands are then processed within the periplasm of *T. forsythia* and subsequently MurNAc and PGN-derived disaccharides are internalized via the inner membrane transporters TfMurT and TfAmpG, respectively [Hottmann, 2021 #19871]. The unique activity of TfNamZ1 fits perfectly with its proposed function in *T. forsythia*. The Tf*namZ1* gene is located upstream of the Tf*ampG* gene, coding for the inner membrane transporter TfAmpG, which we have recently shown to import exogenous PGN-derived GlcNAc-MurNAc and GlcNAc-anhydro-*N*-acetylmuramic acid (anhMurNAc) [Mayer, 2020 #18932]. Thus, the products of TfNamZ1 digestion, disaccharide GlcNAc-MurNAc, can be taken up by the inner membrane transporter TfAmpG, whereas MurNAc, the product of TfNamZ2 (and to some extent also of TfNamZ1), is taken up by TfMurT [Ruscitto, 2016 #14240].

Although we do not have a direct proof that the TfNamZ enzymes function within the periplasm, the N-terminal amino acid sequences of all three proteins contain a putative signal peptide sequence (Sec translocase/signal peptidase I (Sec/SPI) signal sequences) which was confidently identified using the prediction program SignalP-5.0 [Almagro Armenteros, 2019 #15857]. First attempts to purify the three TfNamZs as full length C-terminally His_6_ tagged recombinant proteins in a soluble form failed, presumably because the signal peptides interfered with the expression in *E. coli* as soluble cytoplasmic proteins. However, when we removed the predicted signal peptides, two of the recombinant proteins, TfNamZ1 and TfNamZ2, were obtained in a soluble form in high yield. Interestingly, the predicted cleavage sites for the signal peptides of the TfNamZs are located between an alanine (A) and a glutamine (Q) (see also Fig. S1: A_21_Q_22_ for TfNamZ1 and A_24_Q_25_ for TfNamZ2 and TfNamZ3). The AQ signal peptide cleavage sites predicted for all three TfNamZs (as well as for BfNamZ2) follow the so-called Q-rule, which likely constitutes a characteristic signal peptide cleavage site among *Bacteroidetes* species. Concomitant to signal peptide cleavage, the glutamine residue adjacent to the signal peptide anchored at the inner membrane may get cyclized to a pyroglutamate and released into the periplasm [Bochtler, 2018 #19861].

For TfNamZ3 (Tanf_00855), we failed to obtain a pure soluble protein for so far unknown reasons. This was unexpected, since TfNamZ2 was perfectly soluble and it shares 95% overall identity and a nearly identical signal-peptide with TfNamZ3 (Fig. S2). Thus, further efforts are required to obtain TfNamZ3 in a soluble form for characterization.

In summary, this study revealed that members of the glycosidase family 171 can have distinct substrate specificity and activity. They either solely release terminal MurNAc residues from the non-reducing ends of substrates, or they possess a somehow broader specificity and can cleave terminal GlcNAc-MurNAc and GlcN-MurNAc disaccharides from the non-reducing ends. Both activities are present in two members of GH171 from *T. forsythia* and may allow this oral pathogen, which is unable to *de novo* synthesize the MurNAc content of its own PGN, to acquire MurNAc and disaccharides containing MurNAc from surrounding bacteria within its oral habitat.

## MATERIALS AND METHODS

### Chemicals, enzymes, and oligonucleotides

Enzymes for DNA cleavage and cloning were purchased from New England Biolabs (Ipswich, MA) or Fisher Scientific (Schwerte, Germany). The Gene JET plasmid miniprep kit, PCR purification kit, Gene Ruler 1-kb marker and isopropyl β-D-thiogalactopyranoside (IPTG) were obtained from Fisher Scientific. 2-Acetamido-4-O-(2-acetamido-2-deoxy-β-D-glucopyranosyl)-2-deoxy-D-muramic acid (GlcNAc-MurNAc) and 4-nitrophenyl *N*-acetyl-β-D-glucosaminide (pNP-GlcNAc) were obtained from Carbosynth/Biozol Diagnostica (Eching, Germany). pNP-MurNAc was synthesized as previously described [Müller, 2021 #18933]. Lysogeny broth (LB Lennox) medium, kanamycin sulfate and acetonitrile (ROTISOLV^®^ HPLC ultra gradient grade) were obtained from Carl Roth GmbH (Karlsruhe, Germany). Oligonucleotide primers, which are listed in Table S1, were purchased from MWG Eurofins (Ebersberg, Germany) or Sigma-Aldrich (St. Louis, USA). Formic acid and ammonium formate for LC-MS analysis were purchased from Sigma-Aldrich and Merck (Darmstadt, Germany), respectively.

### Bacterial strains, plasmids, and growth conditions

The bacterial strains and plasmids used in this study are presented in Table S2. *Escherichia coli* DH5α and BL21(DE3) cells were grown in LB Lennox medium under continuous 150 rpm shaking at 37 °C. The LB medium was solidified with 1.5% (w/v) agar, if required*. E. coli* cells containing pET29b or pET28a derived plasmids were cultured in LB medium supplemented with kanamycin (50 µg/ml) and pET16b derived plasmids were maintained by adding ampicillin (100 µg/ml).

### Cloning, expression and purification of recombinant proteins

Recombinant BsNagZ, BsNamZ, BsCwlC, SaAtl^AM^ and SaAtl^Glc^ enzymes were purified as previously described [Litzinger, 2010 #12434;Büttner, 2014 #13793;Walter, 2021 #16248;Müller, 2021 #18933]. For the cloning of *TfnamZ1- 3* the respective genes were amplified by PCR using primer pairs as listed in Table S1 and genomic DNA from *T. forsythia* strain ATCC 43037 (NCBI, Reference Sequence: NZ_JUET00000000.1) [Friedrich, 2015 #15480;Zwickl, 2020 #19875]. Importantly, the previously deposited genome in NCBI with the GenBank accession number CP00319 [Dewhirst F., Tanner A., Izard J., Brinkac L., Durkin A.S., Hostetler J., Shetty J., Torralba M., Gill S., Nelson K, Submitted (12-DEC-2011) The J. Craig Venter Institute, 9704 Medical Center Dr., Rockville, MD 20850, USA], as well as entries in the Uniprot database (www.uniprot.org) correspond to the *T. forsythia* strain 9A2A and not to strain ATCC 43037 as annotated (for an explanation, see [Friedrich, 2015 #15480;Zwickl, 2020 #19875]). For the overexpression of the TfNamZs, 1 L LB medium supplemented with 50 μg/ml kanamycin was inoculated with overnight cultures of *E. coli* BL21(DE3) harboring the expression plasmids pET29b-*BsnamZ*, pET29b-*TfnamZ1*, pET29b-*TfnamZ2 or* pET29b-*TfnamZ3* to an initial OD_600_ of 0.05. Cells were grown to log phase (at OD_600_ 0.5 to 0.7) and after cooling the culture medium to RT, expression of the recombinant proteins was induced by addition of IPTG to a final concentration of 1 mM. Bacteria were grown under continuous shaking at 18 °C and harvested (4000*g* for 30 min at 4 °C) after overnight incubation. Bacterial pellets were resuspended in 20 ml of sodium phosphate buffer A (20 mM Na_2_HPO_4_ x 2 H_2_O, 500 mM NaCl with a protease inhibitor cocktail (Roche) each and cells were disrupted using a French Press (Sim-Aminco Spectronic Instruments, Inc) three times at 1000 psi. Sodium phosphate buffer A with pH 7.5 was used for purification of the TfNamZ recombinant enzymes. Cell debris and unbroken cells were removed by centrifugation at 14000 rpm for 60 min at 4 °C. Purification of the C-terminal His_6_-tagged TfNamZ proteins was performed by Ni^2+^ affinity chromatography. Therefore, the obtained supernatants were filtered through sterile PVDF filters (pore size 0.22 μm) and loaded on 1 ml His-Trap columns (GE Healthcare), pre-equilibrated with ten column volumes of each Millipore water and sodium phosphate buffer A, using a protein purification system (ÄKTApurifier, GE Healthcare). Elution of the proteins was achieved by using a linear gradient of imidazole from 4 mM to 500 mM with sodium phosphate buffer B (20 mM Na_2_HPO_4_ x 2H_2_O, 500 mM NaCl, 500 mM imidazole, pH 7.5). Purity of the *T. forsythia* enzymes was analyzed by 12% SDS-PAGE after staining with Coomassie Brilliant Blue G250 dye. Peak fractions containing desired proteins were pooled and further purified by SEC (HiLoad 16/60 Superdex 200 column, GE Healthcare) using sodium phosphate buffer A with pH 7.5 as eluent. Peak fractions were analyzed for pure proteins by 12% SDS-PAGE and fractions containing pure enzymes were pooled and concentrated using Vivaspin concentrators with 30 kDa cut-off filter (Sartorius). Protein concentrations were determined using the corresponding extinction coefficient at 280 nm (ExPASy, ProtParam tool) and measured in a 1 ml quartz cuvette (Hellma Analytics) using a SpectraMax M2 spectrophotometer (Molecular Devices). Proteins were long-term stored at -80 °C in 10% glycerol.

### PGN isolation from *B. subtilis* 168 and *S. aureus* USA300 JE2

For the isolation of PGN from *B. subtilis* strain 168, 1 L LB medium was inoculated with an overnight culture to an initial OD_600_ of 0.1. Cells were grown aerobically to an OD_600_ of 1, centrifuged down and resuspended in 15 ml of Tris buffer (100 mM, pH 8) supplemented with 400 µg of proteinase K to prevent self-hydrolysis of PGN by endogenous autolytic enzymes. Afterwards, the bacteria were added dropwise to an equal amount of boiling Tris buffer and boiling was continued under constant stirring for 1hr. The cell pellet was centrifuged (4,000 × g) for 15 min and the pellet was stored at 4 °C. Frozen bacterial cells were resuspended in 6 ml of Tris buffer, containing 600 µg of α-amylase, 100 µg of RNase A and 50 U of DNase I and incubated for 2 hrs at 37 °C under constant shaking. Afterwards, proteinase K (400 µg) was added and the mixture was boiled for 1 hr in an equal volume of SDS (final concentration of 2%). The crude PGN sample was subsequently transferred to 15 ml conical tubes (3148-0050, Thermo Scientific) and sacculi were spun down at 22,000 x g at 40 °C for 30 min in a Heraeus Multifuge X1R using a Fiberlite™ F15-8 x 50cy Fixed Angle Rotor from Thermo Scientific. SDS was removed by washing the sacculi with 60 °C pre-warmed Millipore water several times until no SDS was detected by the methylene blue assay [Hayashi, 1975 #12401]. The wall teichoic acids, which are covalently bound to the MurNAc-moiety of the PGN, were further removed by incubation of the *B. subtilis* sacculi in 5 ml of 1 M HCl at 37 °C for 4 hrs under constant rotation. Afterwards, the PGN sacculi were centrifuged in a table centrifuge (max. speed for 10 min) and the pellet was washed with Millipore water until pH > 6 with pH indicator strips was reached. The pellet was then dried under vacuum at 40 °C and pellets were stored at 4°C.

For the isolation of PGN from *S. aureus* strain USA300 JE2, a previously described protocol [Turner, 2016 #14928] with modifications was applied. Briefly, overnight culture of *S. aureus* was inoculated in 1 L LB medium to an initial OD_600_ of 0.05. Cells were grown for about 20 h aerobically to a OD_600_ of 6.1, centrifuged and pellets were frozen at -80 °C. Furthermore, the bacteria were placed on ice, resuspended in 60 ml ice-cold Millipore water and added dropwise to 60 ml pre-boiled 10% SDS (final concentration of 5%). After 1 hr of constant boiling under stirring, SDS was removed from the sacculi by several washing steps with pre-warmed 60 °C Millipore water until no SDS was detected [Hayashi, 1975 #12401]. The bacterial pellet was resuspended in 12 ml Millipore water and cells were added to glass beads (0.1 mm from Roth) in 15 ml tubes (Precellys). Cells were broken in Precellys Evolution cell disrupter (speed 6.5, four times each for 40 s) and the glass beads were separated from the sacculi by using a 40 µm nylon mesh cell strainer (Fisher Scientific). Obtained *S. aureus* sacculi were collected in 40 ml Millipore water, centrifuged at high speed and redissolved in 20 ml of Tris-HCl (50 mM, pH 7.6) with 10 mM MgCl_2_ and 1200U DNase I and 1000 µg RNase A. Digestions were performed at 37 °C for 3 hrs under constant rotation, sacculi were centrifuged and redissolved in 20 ml of Tris-HCl buffer (50 mM, pH 7.6) with 2 mg Trypsin (Sigma-Aldrich, T-4799). Trypsin digestion was performed overnight at 37 °C under constant shaking. To remove the wall teichoic acid, sacculi were resuspended in 10 ml of 48% hydrofluoric acid for 48 hrs at 4 °C under constant shaking. *S. aureus* sacculi were transferred to Eppendorf tubes, centrifuged in a table centrifuge (13000 rpm for 7 min) and the PGN pellets were washed several times with Millipore water until pH of about 6.5 with pH indicator strips was reached. PGN was dried in a vacuum concentrator and pellets were stored at 4 °C.

### Specificity of TfNamZs to chromogenic pNP substrates

150 µM of the chromogenic substrates pNP-GlcNAc or pNP-MurNAc were incubated with 2 µM of BsNagZ, BsNamZ or TfNamZs enzymes at 37 °C in 50 mM phosphate buffer for pNP-GlcNAc (pH 7) and for pNP-MurNAc (pH 8). Enzymatic reactions with the chromogenic substrates were performed in a round bottom 96 well/plate (Greiner Bio-One) in a final volume of 100 µl. After overnight incubations, reactions were stopped by adding 50 µl of 100 mM carbonate buffer (Na_2_CO_3_/NaHCO_3_, pH 10.8) and pictures were taken to visualize the development of a yellow colour, obtained by the release of pNP in the enzymatic reactions.

### Specificity of TfNamZs for MurNAc-GlcNAc and GlcNAc-MurNAc

Substrate specificity of TfNamZs was also tested with commercially available disaccharide GlcNAc-MurNAc and the disaccharide MurNAc-GlcNAc, which was prepared by enzymatic digestion of isolated PGN sacculi. Therefore, 0.1 mg PGN from *S. aureus* USA300 (prepared as described above) was digested overnight with 5 µg SaAtl^AM^ and 5µg SaAtl^Glc^ enzymes in a final volume of 50 µl in 50 mM phosphate buffer, pH 7.5. The sample was boiled for 10 min at 95 °C and after centrifugation (13,000 rpm; 10 min), the supernatant was further digested with the exo-*N*-acetylglucosaminidase BsNagZ (2.5 µM) for 3hrs (final volume of 50 µl) to cleave the GlcNAc-MurNAc-GlcNAc content of the MurNAc-GlcNAc sample, that was obtained during the digestion of the PGN glycans with the *Sa*Atl^Glc^. After boiling the samples, no enzyme (buffer) or recombinant BsNamZ, TfNamZ1 or TfNamZ2 (each 2.5 µM) enzymes were added to the enzyme reaction and samples were incubated overnight in final volume of 50 µl. Finally, samples were boiled, centrifuged at maximum speed on the table centrifuge and supernatants were stored at 4 °C.

In addition, GlcNAc-MurNAc (0.5 mM) was incubated with either of the recombinant NamZ enzymes from *T. forsythia* or from BsNagZ (2.5 µM each) in phosphate buffer, pH 7 in a final volume of 50 µl. Overnight reactions were stopped by heating the samples at 95 °C for 10 minutes. Samples were then centrifuged (13,000 rpm; 10 min) and the supernatants were kept at 4 °C.

Supernatants were analyzed by LC-MS for the presence of MurNAc-GlcNAc, MurNAc and GlcNAc. The exact monoisotopic masses [M] of the proton adduct [M+H]^+^ of the investigated peptidoglycan sugar metabolites MurNAc-GlcNAc, MurNAc and GlcNAc are summarized in Table S3. The relative metabolite concentrations (% area under the curve, AUC) represent the mean from three biological replicates. The areas under the curve of the respective extracted ion chromatograms were calculated and are presented in %.

### Specificity of TfNamZs for PGN and PGN-derived glycan substrates

PGN from *B. subtilis* (0.5 mg) was resuspended in phosphate buffer (pH 7.0, 50 mM) and incubated with 2.5 µM TfNamZ1 or TfNamZ2. The reaction was incubated at 37 °C overnight under constant rotation and stopped at 95 °C for 10 minutes. The samples were then centrifuged and the supernatants were analyzed by MS.

To generate peptide-free PGN glycan strands from *B. subtilis*, 2 mg PGN was solved in 200 µl Tris-HCl buffer (pH 8, 100 mM) and 5 µg of the MurNAc-L-Ala amidase CwlC from *B. subtilis* (BsCwlC) was added to the murein sacculi. Peptides were digested at 60 °C for 6 hrs, and enzyme reactions were stopped by inactivation at 95 °C for 10 min. ¼ of the CwlC-digested PGN was dried and then solved in a phosphate buffer (pH 7, 100 mM) and incubated overnight with buffer (control) or with 2.5 µM TfNamZ1 or TfNamZ2 enzymes in a final volume of 50 µl. Samples were heat inactivated and the supernatants were analyzed by HPLC-MS.

For the generation of PGN peptide-free glycan strands from *S. aureus*, 0.1 mg of the purified sacculi were incubated with 5 µg of SaAtl^AM^ recombinant enzyme in phosphate buffer (pH 7.5) in a final volume of 40 µl. Sample was incubated at 37°C under constant rotation. After overnight incubation, enzymes were inactivated by heating, samples were centrifuged and 2.5 mM of the TfNamZ or BsNamZ enzymes were added to the supernatants (final volume of 40 µl). After overnight incubation, enzymes were inactivated by heating, centrifuged and the supernatant was analyzed by HPLC-MS.

### LC-MS analysis of disaccharides or PGN-derived substrates

5 µl samples were subjected to LC-MS analysis using an UltiMate 3000 LC system (Dionex) coupled to an electrospray-ionization-time of flight mass spectrometer (MicrO-TOF II; Bruker) that was operated in positive-ion mode. Metabolite separation was performed with a Gemini C_18_ column (150 x 4.6 mm, 110 Å, 5 μm; Phenomenex) at 37 °C with a flow rate of 0.2 ml/min using a 45-min gradient program as described in [Gisin, 2013] with optimized column re-equilibration step: 5 min, 100% buffer A (0.1% formic acid with 0.05% ammonium formate in Millipore water), then 30 min of a linear gradient to 40% buffer B (acetonitrile), and 10 min of 100% buffer A for column re-equilibration. The mass spectra of the samples were analysed with Data Analysis (Bruker) and Prism 8 (GraphPad) software and are shown as base peak chromatograms (BPCs) and extracted ion chromatograms (EICs) of metabolites presented as measured *m/z* values. Theoretical m/z values of the investigated sugar metabolites are summarized in Table S3.

## ACKNOWLEDGMENTS

We thank Prof. Thilo Stehle for providing us with the Atl^AM^ enzyme, Maraike Müller and Isabel Hottmann help in the initial phase of the project and Axel Walter for bioinformatic support. Dirk Hauck (HIPS) is acknowledged for excellent technical support.

## SUPPORTING INFORMATION

This article contains supporting information.

## AUTHOR CONTRIBUTIONS

KB and MB conducted experiments, MB and CM wrote and designed the manuscript, AL and AT conceived ideas and AT provided pNP-MurNAc. All authors read and approved the final version of the manuscript.

## ETHICAL STATEMENT

Ethics approval was not required, as the work includes only an *in vitro* characterization of carbohydrate metabolic enzymes.

## SUPPLEMENTAL MATERIAL

**FIG. S1.**
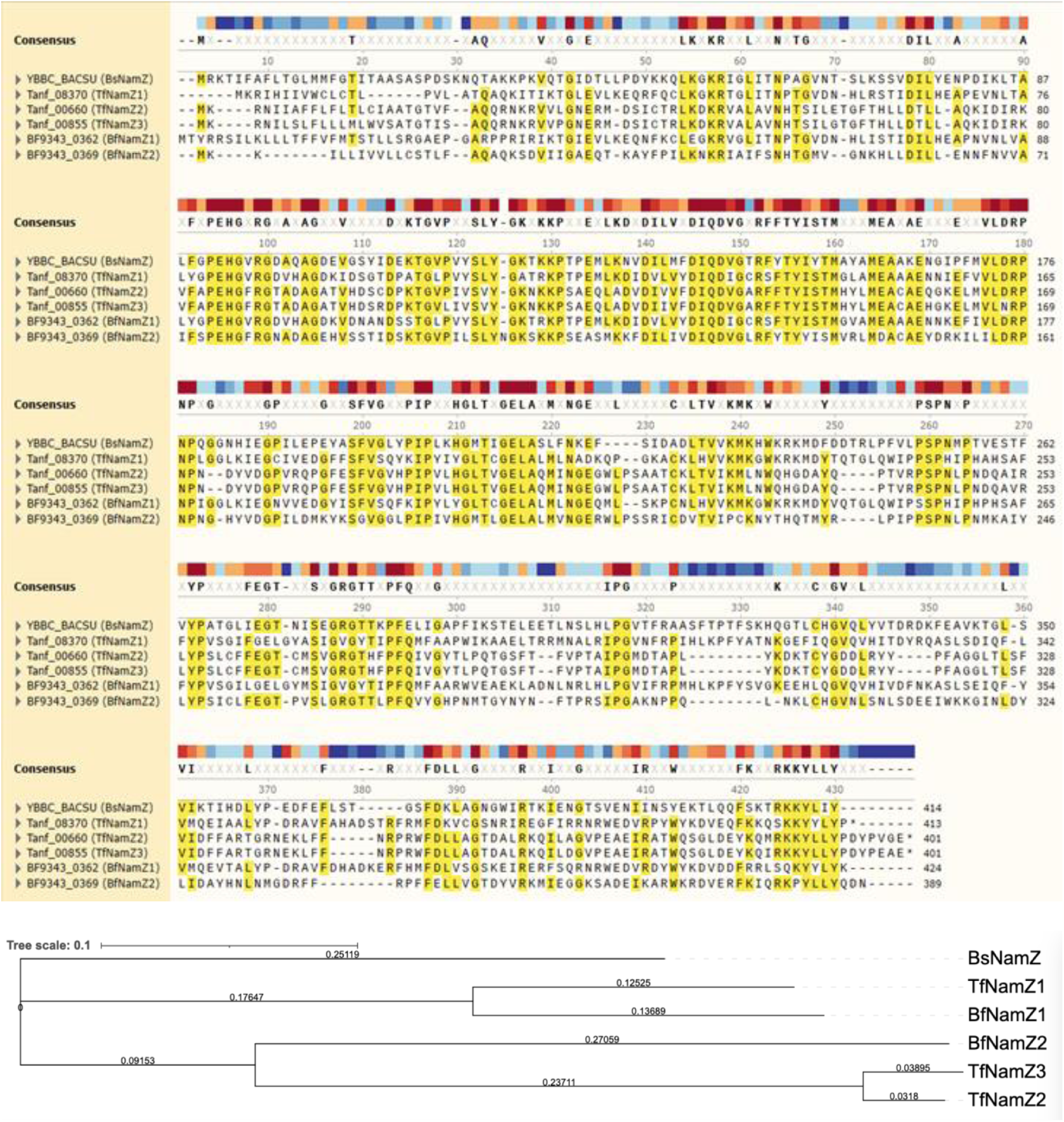
Multiple amino acid sequence alignment and neighbor-joining tree of NamZ orthologs from *B. subtilis* 168 (BsNamZ), *B. fragilis* ATCC 9343 (BfNamZ1 and BfNamZ2) and *T. forsythia* ATCC 43037 (TfNamZ1, TfNamZ2, TFNamZ3). The alignment was conducted with Clustal Omega and was visualized with the SnapGene multiple alignment tool. The phylogenetic, neighbor-joining tree was constructed with iTOL version 6.4 [Letunic, 2021 #20026])

**FIG. S2.**
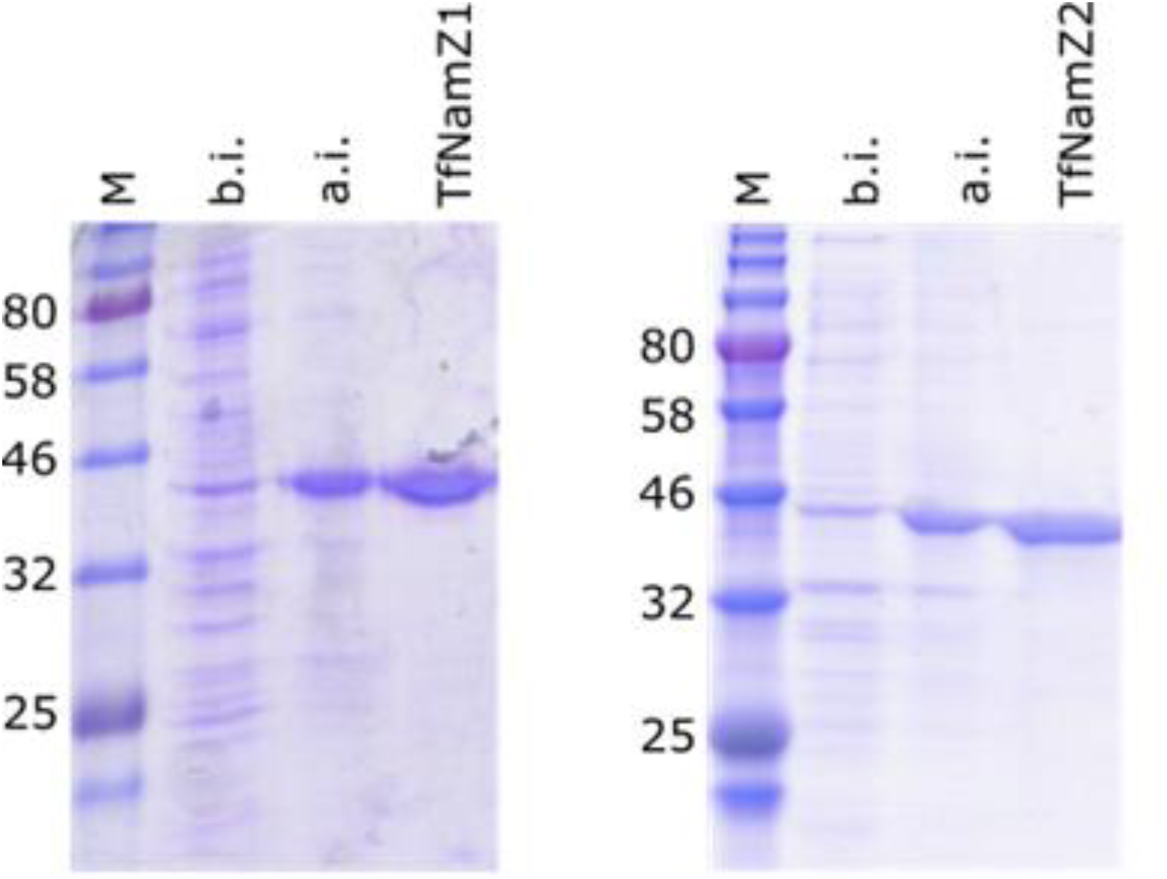
SDS-PAGE analysis of TfNamZs. Expression and purity of the recombinant proteins TfNamZ1 and TfNamZ2 were analyzed by SDS-PAGE with Coomassie brilliant blue staining. Lane 1, protein molecular weight marker (M, size in kDa); lane 2, *E. coli* cell extract before IPTG induction (b.i.), lane 3, after IPTG induction and overnight incubation at 18 °C (a.i.); lane 4, 5 µg of TfNamZ recombinant proteins after purification by Ni^2+^-affinity chromatography and size exclusion chromatography.

**FIG. S3.**
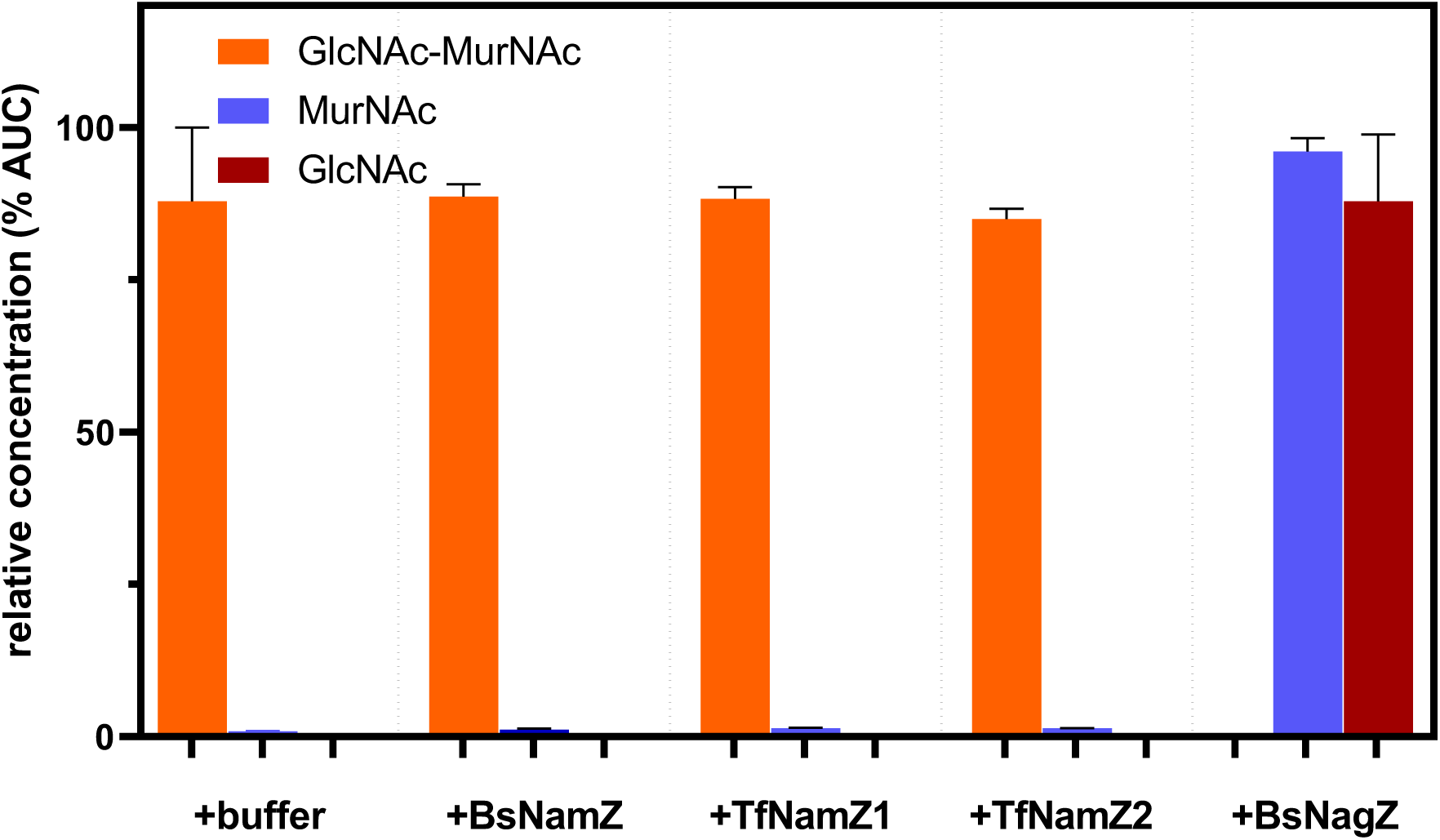
Commercial GlcNAc-MurNAc was incubated with buffer, BsNagZ (positive control), BsNamZ, TfNamZ1 or TfNamZ2 enzymes. Overnight reactions were heat inactivated and the supernatants were analyzed for the presence of GlcNAc-MurNAc (orange), MurNAc (blue) and GlcNAc (red) by HPLC-MS analysis. The exact monoisotopic masses [M] of the proton adduct [M+H]^+^ of the investigated peptidoglycan sugar metabolites MurNAc-GlcNAc, MurNAc and GlcNAc are summarized in Table S3. Data from three biological replicates are presented as areas under the curve from the extracted ion chromatograms (EICs) of the respective compounds and are presented in %. GlcNAc-MurNAc was completely degraded by BsNagZ but, as expected, BsNamZ as well as the TfNamZs enzymes failed to show β-*N*-acetylglucosaminidase activity.

**FIG. S4.**
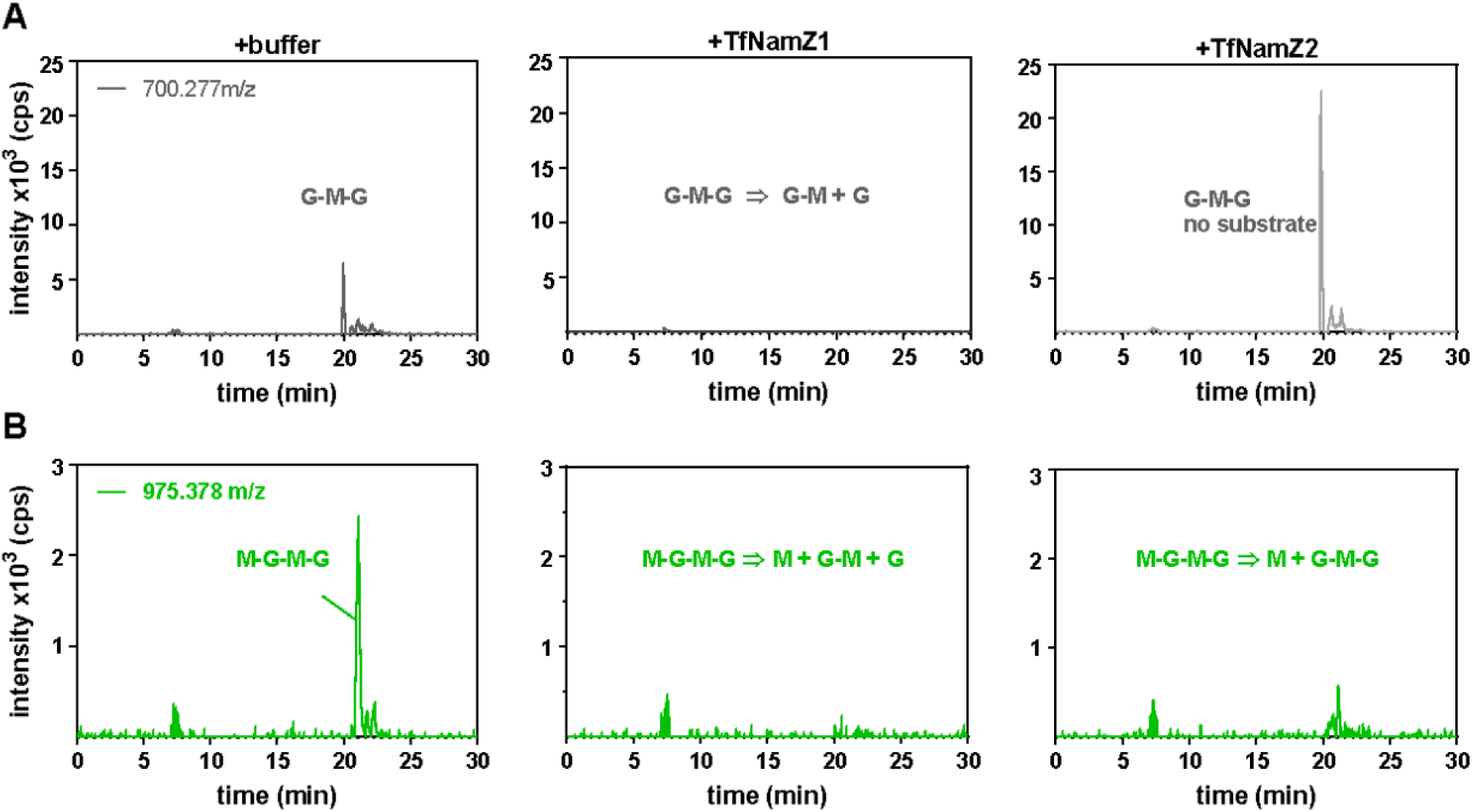
TfNamZ1 and TfNamZ2 show differences in substrate specificity with oligomeric PGN. The peptide-free PGN glycan mixture from *S. aureus*, generated by Atl-digestion as described in Figure 4, contains minor amounts of the trisaccharide **GlcNAc-MurNAc-GlcNAc (G-M-G; A)** and the tetrasaccharides **MurNAc-GlcNAc-MurNAc-GlcNAc (M-G-M-G; B)**, besides the main product MurNAc-GlcNAc. Incubation with the enzymes revealed that TfNamZ1 cleaves both oligosaccharides, whereas TfNamZ2 only cleaves M-G-M-G. While TfNamZ2 cleaves only the MurNAc-moiety from the non-reducing end of M-G-M-G, yielding G-M-G, this product is further cleaved by TfNamZ1, yielding **MurNAc-GlcNAc (**M-G) and **GlcNAc (**G) (as indicated). Data are presented as EICs for the trisaccharide GlcNAc-MurNAc-GlcNAc, G-M-G in grey, ([M+H]^+^ = 700.277 m/z) and for the tetrasaccharide MurNAc-GlcNAc-MurNAc-GlcNAc in green ([M+H]^+^ = 975.378 m/z), and are shown as intensity x 10^3^ (cps).

**FIG. S5.**
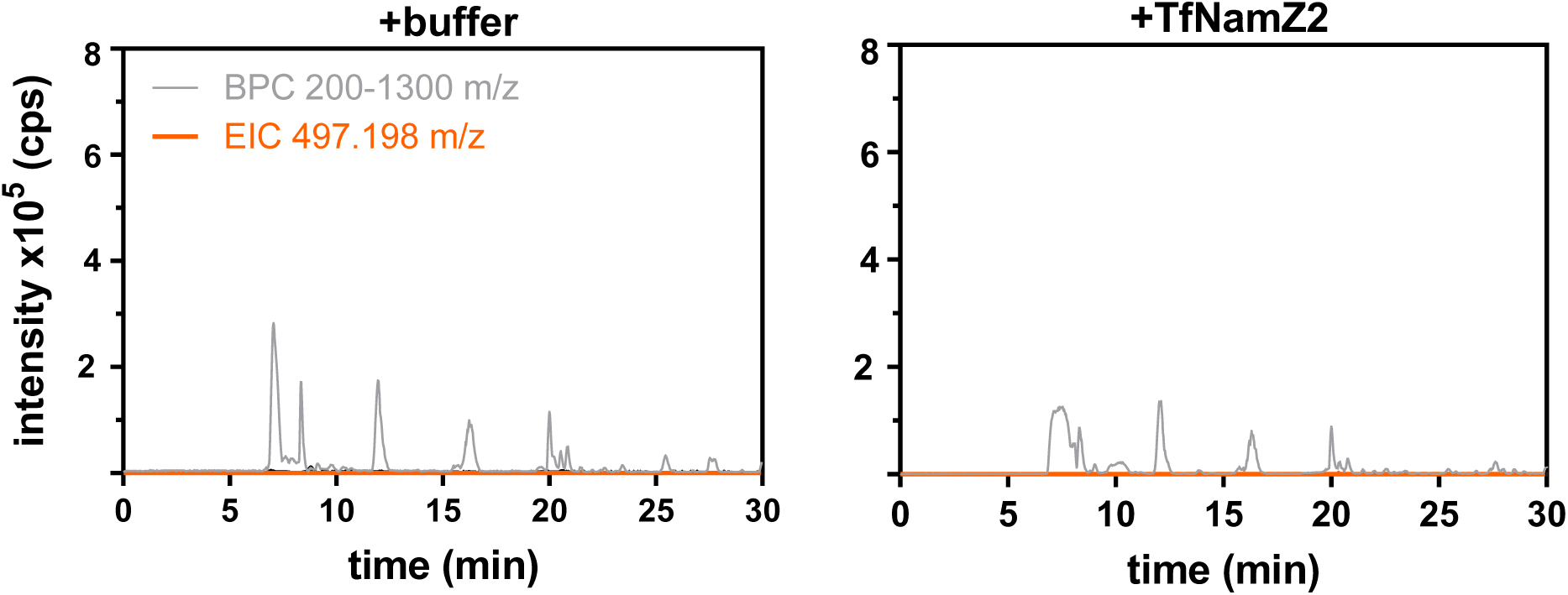
*B. subtilis* glycan strands are not cleaved by TfNamZ2. *B. subtilis* glycan strands were generated as described in Figure 5. Data are presented as base peak chromatograms (BPC) 200-1300 m/z in gray and as extracted ion chromatogram (EIC) of [M+H]^+^ = 497.198 m/z, corresponding to GlcNAc-MurNAc (in orange) and are plotted as intensity x 10^5^ counts per second (cps).

**FIG. S6.**
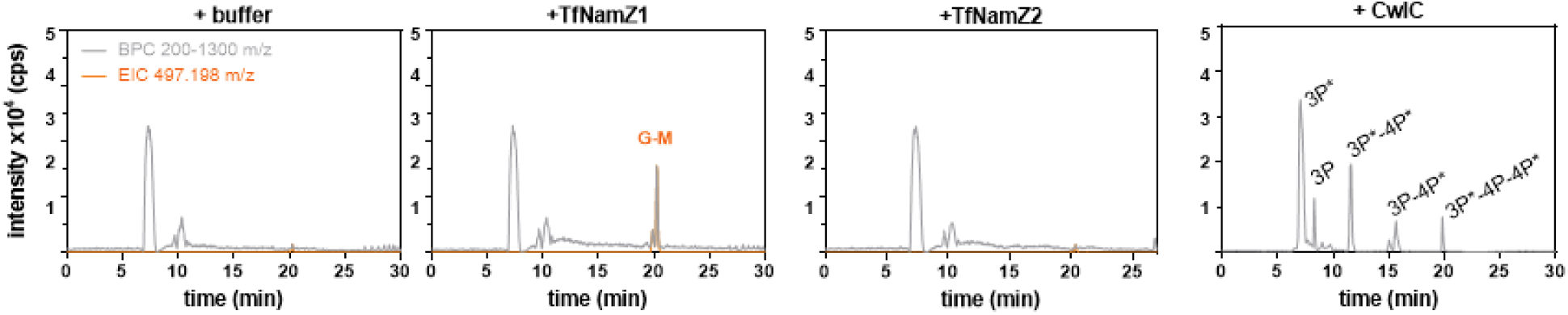
Substrate specificity of TfNamZ1 and TfNamZ2 with *B. subtilis* PGN sacculi. PGN sacculi from *B. subtilis* were incubated with TfNamZ1, TfNamZ2, the amidase BsCwlC, or buffer as a control. Samples were analyzed by HPLC-MS and the data are presented as base BPCss 200-1300 m/z (gray lines) and as extracted ion chromatogram (EIC) for GlcNAc-MurNAc (orange, [M+H]^+^ = 497.198 m/z). The peptide products after CwlC digestion were identified by LC-MS according to their exact masses as: 3P-tripeptide, 3P*-tripeptide amidated; 4P-tetrapeptide, 4P*-tetrapeptide amidated.

## Supplementary Tables

**Table S1.**
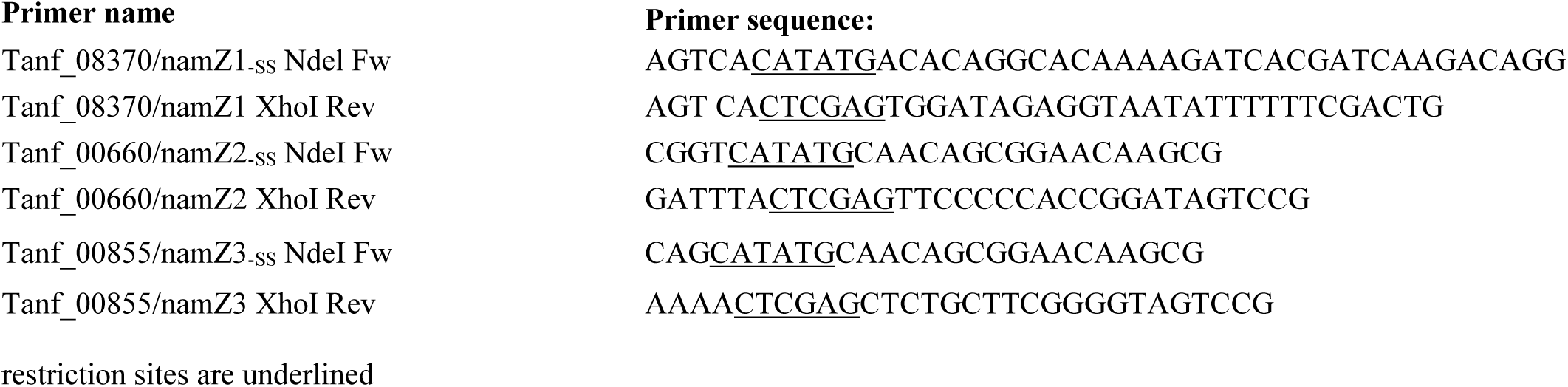
Oligonucleotides used in this study

**Table S2.**
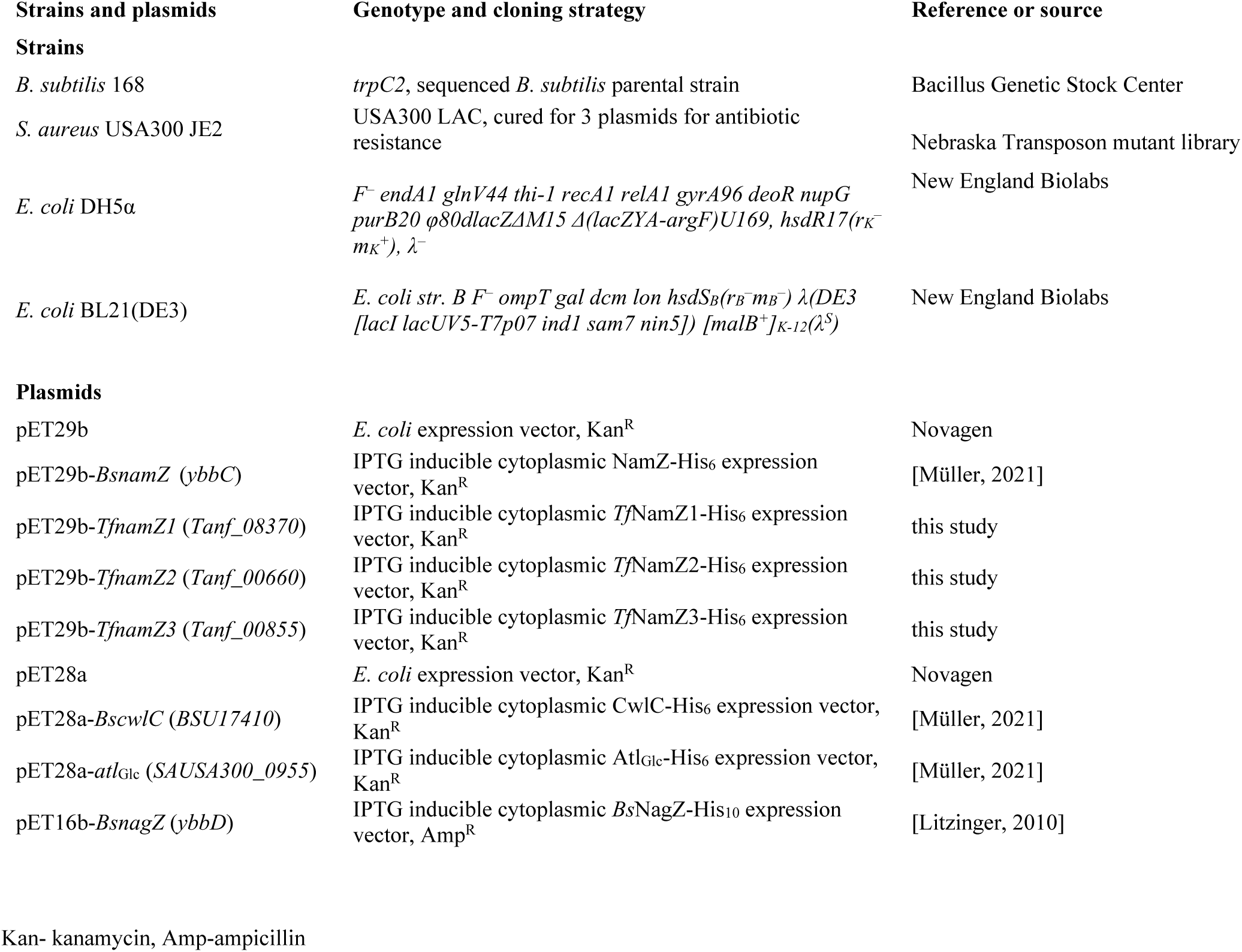
Strains and plasmids used in this study

**Table S3.**
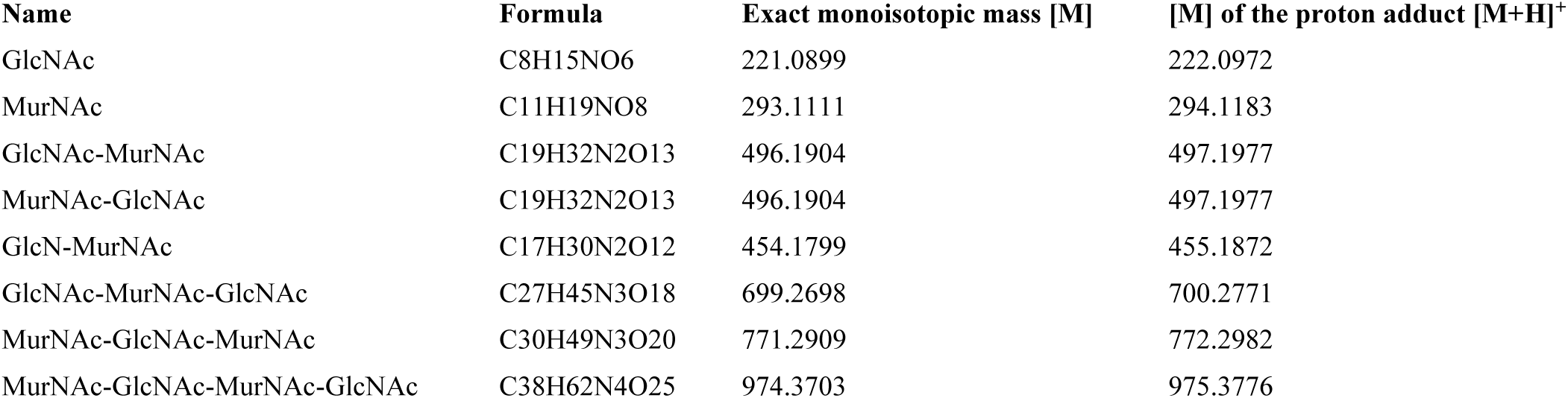
Exact monoisotopic mass [M] and the monoisotopic mass of the proton adduct [M+H]^+^ of relevant PGN carbohydrates

| Name | Formula | Exact monoisotopic mass [M] | [M] of the proton adduct [M+H]<sup>+</sup> |
| --- | --- | --- | --- |
| GlcNAc | C<sub>8</sub>H<sub>15</sub>NO<sub>6</sub> | 221.0899 | 222.0972 |
| MurNAc | C<sub>11</sub>H<sub>19</sub>NO<sub>8</sub> | 293.1111 | 294.1183 |
| GlcNAc-MurNAc | C<sub>19</sub>H<sub>32</sub>N<sub>2</sub>O<sub>13</sub> | 496.1904 | 497.1977 |
| MurNAc-GlcNAc | C<sub>19</sub>H<sub>32</sub>N<sub>2</sub>O<sub>13</sub> | 496.1904 | 497.1977 |
| GlcN-MurNAc | C<sub>17</sub>H<sub>30</sub>N<sub>2</sub>O<sub>12</sub> | 454.1799 | 455.1872 |
| GlcNAc-MurNAc-GlcNAc | C<sub>27</sub>H<sub>45</sub>N<sub>3</sub>O<sub>18</sub> | 699.2698 | 700.2771 |
| MurNAc-GlcNAc-MurNAc | C<sub>30</sub>H<sub>49</sub>N<sub>3</sub>O<sub>20</sub> | 771.2909 | 772.2982 |
| MurNAc-GlcNAc-MurNAc-GlcNAc | C<sub>38</sub>H<sub>62</sub>N<sub>4</sub>O<sub>25</sub> | 974.3703 | 975.3776 |

